# Tau aggregate replication occurs at the pre-synapse of cultured human neurons and increases with application of TNFɑ

**DOI:** 10.64898/2026.05.30.728997

**Authors:** Shekhar Kedia, Emre Fertan, Arkoprovo Paul, Valentina Davi, George Nolan, Koby Baranes, Yunzhao Wu, Jae Eun Kim, Annelies Quaegebeur, Mark R. N. Kotter, Edward Avezov, Georg Meisl, David Klenerman

**Author notes:** **Correspondence to: Professor Sir David Klenerman**, Yusuf Hamied Department of Chemistry, University of Cambridge, UK, CB2 1EW. Co-first authors (Shekhar Kedia, Emre Fertan, Arkoprovo Paul). Current address (Emre Fertan).

## Abstract

Tau aggregation at synapses is a key process driving Alzheimer’s disease but the mechanism(s) that cause this have not been established. We used a model system of forward-programming induced glutamatergic neurons (iNeurons) with three independent cell lines treated with TNFɑ. Using aggregate-specific SIMOA, STED microscopy, and SynPull to detect nanoscopic tau aggregates in bulk samples and at individual synapses, we found that TNFɑ-driven tau aggregation occurs preferentially at the pre-synapse, forming predominantly non-fibrillar aggregates that are larger than ones in the extra- and post-synaptic regions. Using mathematical models of aggregate formation, we fitted the frequency of AT8-positive tau aggregates in synaptosomes, which showed that aggregate replication is the dominant process and is much faster than de-novo aggregate formation, leading to rapid local amplification once one aggregate is formed. Our results provide direct evidence for tau aggregate replication at the pre-synapse, linking inflammation induced tau aggregation with synaptic pathology.

**Graphical abstract:** 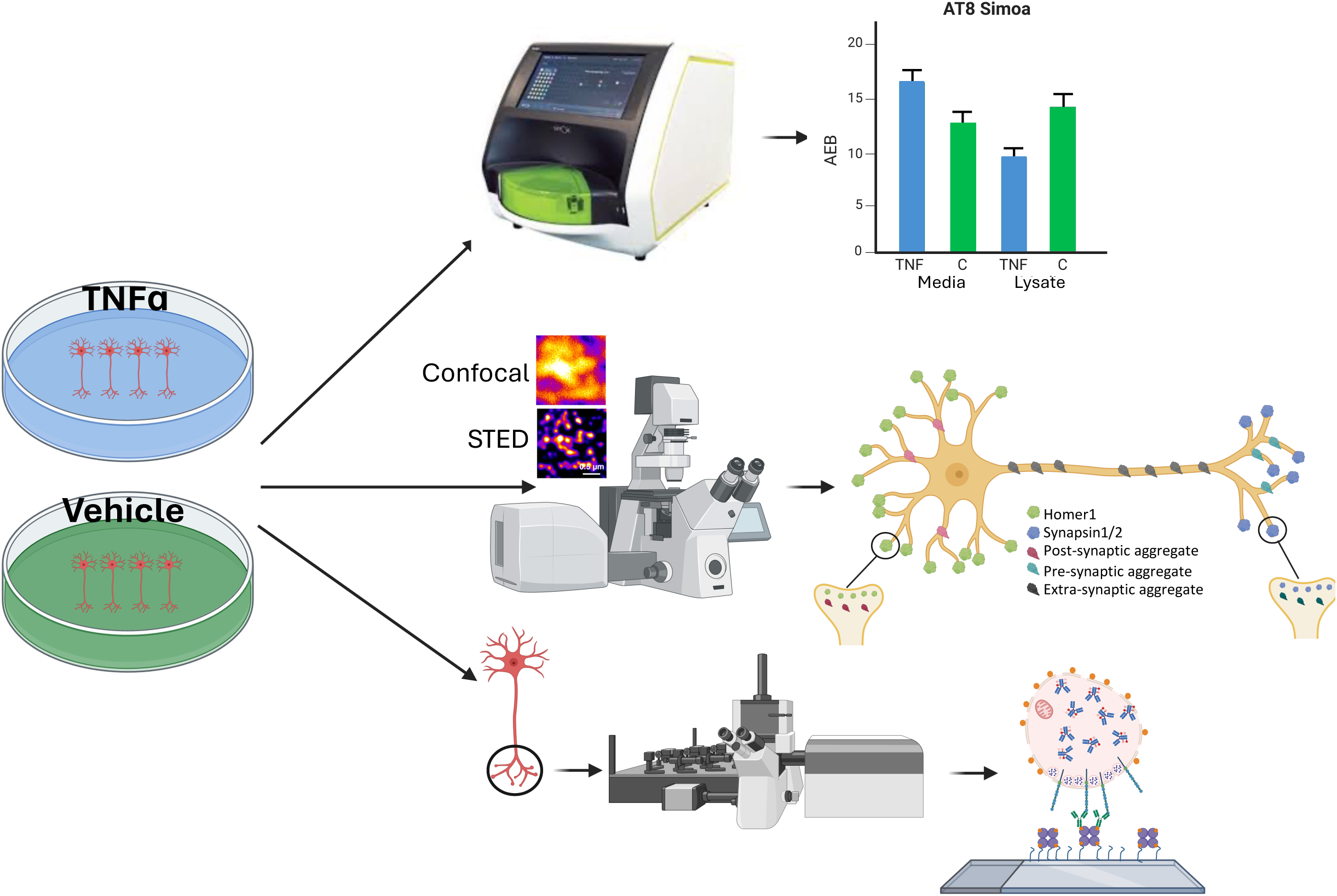

## 1. Introduction

The neuropathological hallmarks of proteinopathies such Alzheimer’s disease (AD)^1^ have traditionally been microscopic, fibrillar, and insoluble aggregates^2^ such as beta-amyloid plaques^3^ and neurofibrillary tau tangles^4^. While the atomic structures of the filaments forming these large aggregates have been identified in elegant studies using cryogenic electron microscopy (cryoEM)^5–7^, basic biophysical principles suggest that these filaments consisting of thousands of individual proteins are highly unlikely to form spontaneously from soluble monomers. Instead, smaller intermediate species will necessarily appear on the pathway to formation of these larger and insoluble aggregates^8^. The presence of these intermediate species -which are often called “oligomers” or “nanoscopic aggregates”, in the brain is sometimes debated^9^, however both theoretical chemistry and experimental findings^10–15^ indicate their presence. Moreover, recent studies have suggested that these oligomeric species are not only present in the brain, but are also responsible for the majority of aggregate induced toxicity^16–21^, meanwhile the larger, insoluble species may be actively produced^22,23^ and even protective to an extent^16,24^. As such, it is crucial to study these oligomeric aggregates and understand their formation and toxicity mechanisms, to get an understanding of the pathogenesis and pathophysiology of proteinopathies and treat them.

However, the characterisation of oligomeric aggregates remains challenging for several reasons. First, they are very small in size, in the nanoscale range, and heterogeneous in shape. As such, they cannot be resolved using traditional light microscopy techniques, due to the diffraction-limit^25^ and also cannot be studied reliably by cryoEM, as they lack a stable homogeneous structure. Second, their abundance is much lower than monomeric species, making it difficult to quantify them using bulk techniques such as ELISA or Western blot. Third, they can be transient and meta-stable, meaning that great care needs to be taken to not alter their structures or cause them to dissociate before they are measured. Therefore, specialised techniques with high enough sensitivity and resolution are needed to detect and characterise these aggregates. Single-molecule detection methods are suitable for studying these aggregates; for instance, digital-ELISA assays such as single molecule array (SIMOA) can be used to quantify their concentration in liquid samples^26^ and super-resolution microscopy techniques such as DNA point accumulation in nanoscale topography (DNA-PAINT)^27^, direct stochastic optical reconstruction microscopy (*d*STORM)^28^, or stimulated emission depletion microscopy (STED)^29^ can be used to characterise their size and shape. Our group has been developing these single-molecule detection methods to study nanoscopic aggregates in post-mortem disease brain samples, which led to the discoveries that the AD therapeutic lecanemab preferentially binds to smaller beta-amyloid aggregates that are present in early AD^30^, that oligomeric tau aggregates are not shared between tauopathies in terms of their structure and post-translational modifications^31^, and that only a subset of cells show altered alpha-synuclein aggregation with larger aggregates in PD^32^.

Experimental measurements combined with kinetic modelling has demonstrated that aggregation occurs by a nucleation step resulting in the initial formation of an aggregate, an elongation step where the aggregate grows by the addition of monomers, and a replication step whereby an aggregate produces another aggregate by either fragmentation or by surface templated nucleation. This replication can lead to the rapid increase of the number of aggregates present, since it bypasses the slower nucleation step, and may thus play a key role in pathology^33^. *In-vitro* experiments, combined with mechanistic analysis have revealed these processes as the intrinsic steps of protein aggregation^34^. While this establishes the baseline behaviour, understanding aggregate formation *in-vivo* requires consideration of additional factors and processes. We have recently extended our kinetic modelling approach to the situation in a cell where there are also processes of aggregate removal. We find that the cells in the brain can exist in a “healthy state” where the aggregates are removed as fast as they are formed and a “diseased state” where the removal processes are no longer sufficient and there is run-away aggregation, with cell stress being able to push the system from one state to another^35,36^.

Nevertheless, post-mortem brain studies are limited in terms of mechanistic interpretation, since the human brain cannot be manipulated and can only be studied as a snapshot. Therefore, to obtain more granular mechanistic insights, cellular model systems are needed. Using an induced pluripotent stem cell (iPSC)-derived neuronal model, we have previously shown that chronic immune stress with tumour necrosis factor alpha (TNFɑ) can lead to the increased formation of beta-amyloid and tau oligomers, importantly in the absence of any mutations, which are then released to the conditioned-media in a size-dependent manner^37^. The analyses in that study used cell lysates and conditioned media, thus the cellular localisation of these aggregates could not be studied. More recently we have also shown that pathological tau aggregation in the AD brains may start at the pre-synaptic terminals, due to reduced removal or increased aggregation in these locations^38^, agreeing with other findings of synaptic localisation of oligomeric tau^39–41^. Since synaptic dysfunction and loss occur early in AD, it is important to determine the aggregation mechanisms of tau at neuronal synapses in a reproducible model system.

Here we employed the newly developed forward-programming iPSC-derived glutamatergic neuronal models (iNeurons), which show neuronal markers rapidly after differentiation on days *in-vitro* (DIV) day 7, along with synaptic markers and electrophysiological activity^42^. We have treated these iNeurons with TNFɑ at 100 ng/ml for 18 days and used three different single-molecule detection and super-resolution methods to quantify and characterise the oligomeric tau aggregates. First, we used aggregate-specific single-molecule array (SIMOA) to quantify the total AT8-positive (referred as AT8+ from here-onwards) tau aggregate load in the cell lysates and conditioned media. Then, we employed STED microscopy^29,43^ to identify the neuronal localisation of tau aggregates, using antibodies specific to pre- and post-synaptic terminals, along with anti-tau antibodies recognising aggregates at different stages. Lastly, we used SynPull, which is a single-molecule pulldown (SiMPull) and *d*STORM microscopy-based method we have developed, allowing the detailed study of synaptosomes at nanometre resolution^38,44^. By combining these single-molecule and super-resolution detection methods with a stochastic model of aggregate formation and removal, we were able to model pathological tau aggregation and show that aggregate replication is the dominant mechanism at the pre-synaptic terminals, which is further elevated with TNFɑ treatment.

## 2. Results

We have used advanced single-molecule detection and super-resolution imaging techniques that we have developed including aggregate-specific SIMOA^26^, STED^29,43^, and SynPull^38,44^, to characterise the nanoscopic tau aggregates in three independent iNeuron lines treated with TNFɑ for 18 days, along with untreated controls. Results were analysed using linear regression with treatment as the predictor of interest and type if iNeuron (the three independent lines) was included as a fixed-effect covariate to account for variability between the cell-lines. Differences between the samples were determined using 95% confidence intervals^45^ and are presented in **Table 1**.

**Table 1.** Statistical differences between the TNFɑ-treated and control samples from three independent forward-programming induced glutamatergic neuronal lines calculated using 95% confidence intervals.

|  |  |
| --- | --- |
| **<u>Luciferase (cell viability)</u>**<br>ioGlutamatergic Neurons:<br>Kolf2 iPSCs:<br>i3 neurons: | **<u>95% confidence interval</u>**<br>$CI_{95} = -309.20, 201.86$<br>$CI_{95} = -62.88, 200.88$<br>$CI_{95} = -31.47, 144.47$ |
| **<u>STED with MC1 antibody</u>**<br>Total number of aggregates:<br>Extra-synaptic aggregates:<br>Number of pre-synaptic aggregates:<br>Number of post-synaptic aggregates:<br>Extra-synaptic aggregate length:<br>Pre-synaptic aggregate length:<br>Post-synaptic aggregate length:<br>Extra-synaptic aggregate eccentricity:<br>Pre-synaptic aggregate eccentricity:<br>Post-synaptic aggregate eccentricity: | **<u>95% confidence interval</u>**<br>$CI_{95} = -56.46, 72.84$<br>$CI_{95} = -58.91, 75.18$<br>$CI_{95} = -14.28, 16.55$<br>$CI_{95} = -12.96, 10.92$<br>$CI_{95} = 8.34, 21.91$<br>$CI_{95} = -70.41, 18.81$<br>$CI_{95} = -118.53, 57.70$<br>$CI_{95} = 0.000, 0.01$<br>$CI_{95} = 0.01, 0.07$<br>$CI_{95} = -0.01, 0.06$ |
| **<u>STED with AT8 antibody</u>**<br>Total number of aggregates:<br>Number of extra-synaptic aggregates:<br>Number of pre-synaptic aggregates:<br>Number of post-synaptic aggregates:<br>Extra-synaptic aggregate length:<br>Pre-synaptic aggregate length:<br>Post-synaptic aggregate length:<br>Extra-synaptic aggregate eccentricity:<br>Pre-synaptic aggregate eccentricity:<br>Post-synaptic aggregate eccentricity: | **<u>95% confidence interval</u>**<br>$CI_{95} = -56.98, 24.58$<br>$CI_{95} = -52.25, 26.21$<br>$CI_{95} = 0.06, 12.83$<br>$CI_{95} = -7.78, 7.96$<br>$CI_{95} = -22.90, 5.17$<br>$CI_{95} = 18.59, 226.20$<br>$CI_{95} = -18.89, 95.47$<br>$CI_{95} = 0.00, 0.02$<br>$CI_{95} = 0.03, 0.12$<br>$CI_{95} = -0.05, 0.06$ |
| **<u>STED with PHF13.6 antibody</u>**<br>Total number of aggregates:<br>Number of extra-synaptic aggregates:<br>Number of pre-synaptic aggregates:<br>Number of post-synaptic aggregates:<br>Extra-synaptic aggregate length:<br>Pre-synaptic aggregate length:<br>Post-synaptic aggregate length:<br>Extra-synaptic aggregate eccentricity:<br>Pre-synaptic aggregate eccentricity:<br>Post-synaptic aggregate eccentricity: | **<u>95% confidence interval</u>**<br>$CI_{95} = -55.69, 38.47$<br>$CI_{95} = -48.91, 36.49$<br>$CI_{95} = -18.42, 9.98$<br>$CI_{95} = -10.72, 9.57$<br>$CI_{95} = -2.77, 23.98$<br>$CI_{95} = -40.81, 71.86$<br>$CI_{95} = 4.64, 100.36$<br>$CI_{95} = -0.01, 0.01$<br>$CI_{95} = -0.03, 0.04$<br>$CI_{95} = -0.04, 0.05$ |
| **<u>Synaptic morphology (STED)</u>**<br>Pre-synaptic count:<br>Post-synaptic count:<br>Pre-synaptic diameter:<br>Post-synaptic diameter:<br>Pre-synaptic area:<br>Post-synaptic area: | **<u>95% confidence interval</u>**<br>$CI_{95} = 2.92, 55.59$<br>$CI_{95} = -36.80, 5.29$<br>$CI_{95} = 37.00, 52.48$<br>$CI_{95} = -1.30, 11.53$<br>$CI_{95} = 14685.07, 20657.96$<br>$CI_{95} = 857.49, 5196.14$ |
| **<u>SynPull with AT8 antibody</u>**<br>Number of aggregates per synaptosome:<br>Fraction of synaptosomes with aggregates:<br>Aggregate length:<br>Aggregate area:<br>Aggregate eccentricity:<br>Phosphatidylserine fraction:<br>Phosphatidylserine and tau aggregate fraction:<br>CD47 fraction:<br>CD47 and tau aggregate fraction:<br>Synaptogyrin fraction:<br>Synaptogyrin and tau aggregate fraction: | **<u>95% confidence interval</u>**<br>$CI_{95} = 0.54, 1.20$<br>$CI_{95} = -2.89, 13.64$<br>$CI_{95} = 0.33, 20.85$<br>$CI_{95} = -721.74, 281.52$<br>$CI_{95} = -0.02, 0.02$<br>$CI_{95} = -18.50, 11.31$<br>$CI_{95} = -11.49, 25.83$<br>$CI_{95} = -46.90, -0.03$<br>$CI_{95} = -8.46, 13.39$<br>$CI_{95} = 7.60, 42.75$<br>$CI_{95} = -1.68, 18.88$ |

### 2.1 Cell viability

To determine the effect of chronic TNFɑ treatment on the viability of iNeurons, we performed a luciferase assay, which reflects the amount of metabolically active cells in the culture. These values were also used to normalise the measures reported for the SIMOA experiments below. While there was no significant difference between the TNFɑ-treated and control iNeurons for any of the three lines (all p > 0.05), significant differences were observed between the cell lines: the greatest viability was observed in the Kolf2 iPSCs, followed by the ioGlutamatergic Neurons and the greatest amount of cell loss was observed in the i3 neurons (AIC = 391.24, F = 44.05, p < 0.001; **Figure 1a**).

**Figure 1.**
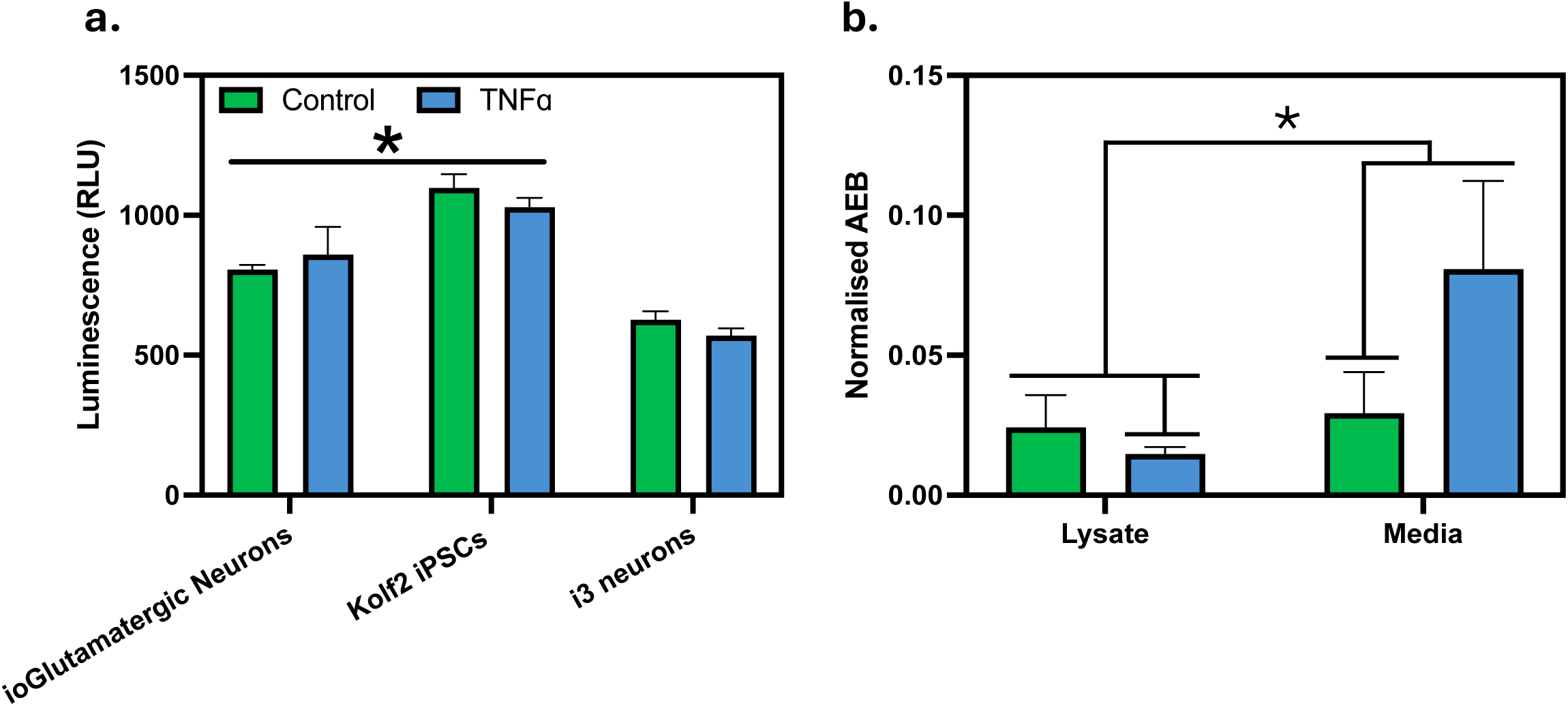
(a) Viability of three independent forward-programming induced glutamatergic neuronal lines was determined using a luciferase assay. (b) AT8-positive tau aggregate levels are measured in the lysates and conditioned media from the iNeurons using aggregate-specific SIMOA, with results normalised to the luciferase assay.

### 2.2 Aggregate specific SIMOA

In-house developed^26^ aggregate specific SIMOA assays were used to quantify the total intra- (cell lysate) and extra-cellular (conditioned media) AT8-positive tau aggregates. There was a significant treatment by sample type (lysate vs media) interaction as the intra-cellular AT8-positive tau aggregate concentration was reduced by TNFɑ treatment, the extra-cellular concentration was increased (AIC = 119.24, F = 12.16, p = 0.002; **Figure 1b)**, indicating that a chronic inflammatory challenge is leading to greater tau aggregate production in iPSC-derived neurons, which are then released from the cells to the extra-cellular space.

### 2.3 STED imaging

While aggregate-specific SIMOA allows the quantification of total aggregate levels in the neuronal lysates and conditioned media, it does not provide any information on the spatial distribution of aggregates inside the iNeurons. Since our previous works have shown that the pathological tau aggregation in the AD brain may begin in the synapses^38^, we complimented SIMOA with STED microscopy, which enables the study of different parts of the neurons. We utilised pre- and post-synaptic markers (synapsin1/2 and homer1, respectively) to compare the nanoscopic tau aggregates between the pre- and post-synaptic compartments along with the extra-synaptic regions such as the neurites (but not the soma). Monoclonal antibodies MC1, AT8, and PHF13.6 were used to study tau aggregates at different aggregation states (aggregates detected by these antibodies are referred to as MC1+, AT8+, and PHF13.6+, respectively, from here onwards). MC1 is a conformation-specific antibody, which recognises discontinuous conformational epitopes involving amino acids 7-9 and 312-322, which are believed to be exposed as an early conformational change, eventually leading to the formation of paired helical fillaments^46^. Meanwhile, the AT8 antibody, which is also used in our SIMOA assay, recognises tau phosphorylated at Ser202 and Thr205, which is subsequent to the conformational change and involved in AD pathogenesis^47^. Lastly, the PHF13.6 antibody recognises tau phosphorylated at Ser396/404, which is considered a later event^48,49^. As such, when used together, these three antibodies can identify tau aggregates at different stages.

The total number of MC1+ tau aggregates did not differ between the treatment conditions; which was further confirmed for the extra-, pre-, and post-synaptic compartments (**Figure 2a**) individually. It should be noted that the extra-synaptic aggregates are characterised by their proximity to the pre- (Synapsin1/2) and post-synaptic (Homer) markers and include the aggregates in the axons and dendrites, respectively, but do not include the ones in the soma (further explained in Methods).

**Figure 2.**
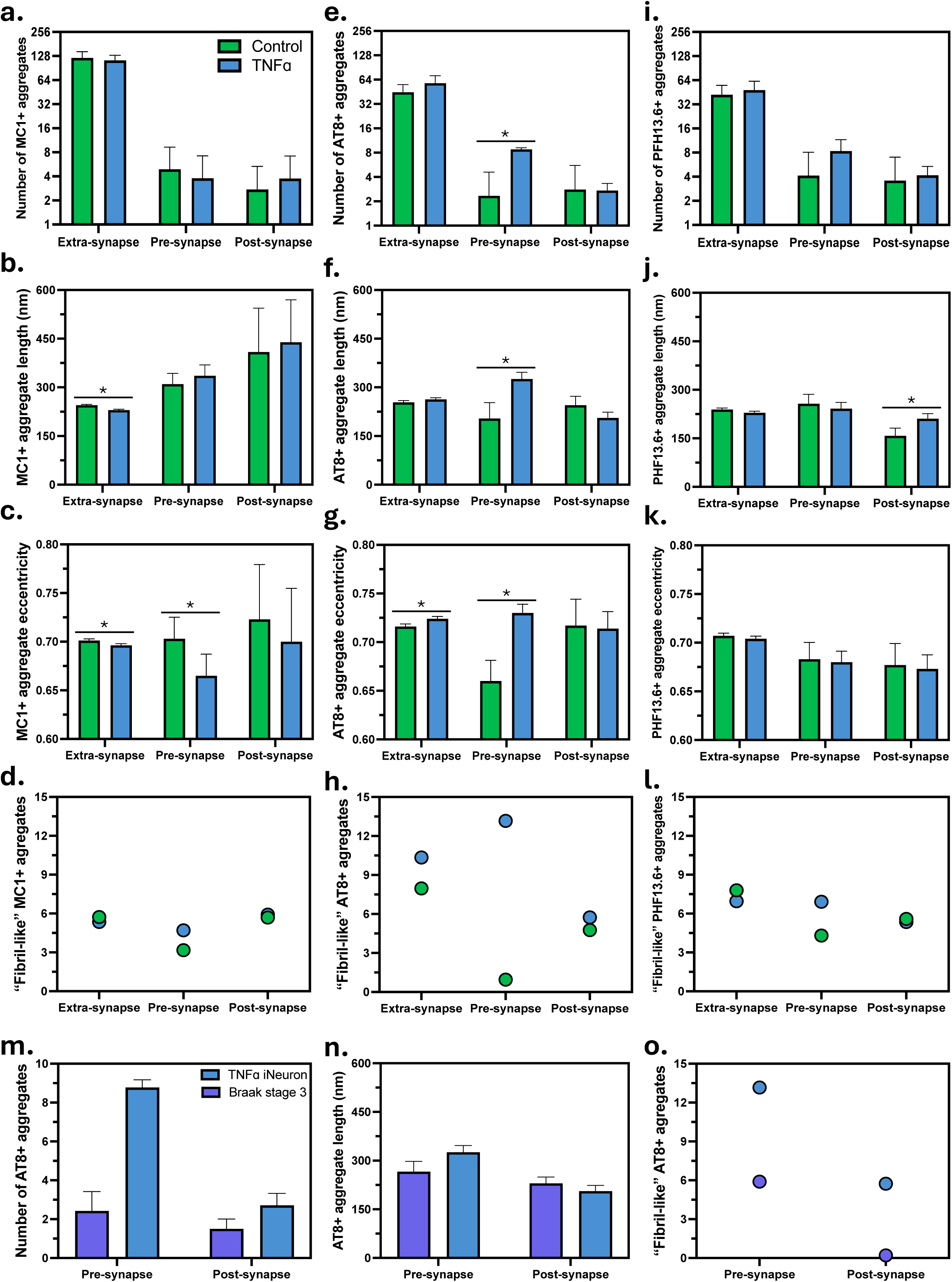
Tau aggregate morphology measured by STED in iNeurons and human AD brains samples. Number (a, e, i), length in nanometres (b, f, j), and eccentricity (c, g, k) of the MC1, AT8, and PFH13.6 positive tau aggregates, respectively. The percentage of fibril-like tau aggregates with an eccentricity value above 0.9 and length over 300 nm for the MC1 (d), AT8 (h), and PFH13.6 (l) antibodies. Number (m), length (n), and shape (o) of AT8-positive tau aggregates in human post-mortem Braak stage 3 pre-frontal cortex samples. Values are plotted as the estimated marginal (least-square) means plus the standard error.

Following their quantification, we further characterised the MC1+ tau aggregates in terms of their length and shape (eccentricity with values close to 0 indicating circular aggregates and values close to 1 indicating flatter, fibril-like aggregates). Length of the MC1+ aggregates was decreased with TNFɑ treatment in the extra-synaptic regions with an average of ∼240 nm in the vehicle treated controls and 230 nm in the TNFɑ treated iNeurons. Meanwhile, aggregate length did not change in the pre- and post-synapse but was overall larger compared to the extra-synaptic regions, with an average of ∼380 nm in the pre-synapse and ∼400 nm in the post-synapse (**Figure 2b**). Eccentricity of the MC1+ aggregates did not differ between the treatments in the post-synaptic region, but the extra- and pre-synaptic aggregates in the TNFɑ-treated iNeurons were less fibrillar than the ones in the vehicle treatment (**Figure 2c**). Lastly, we calculated the proportion of fibrillar aggregates detected with MC1, using the eccentricity and length values; an aggregate with an eccentricity value above 0.9 and length over 300 nm was considered fibrillar. For the extra-synaptic aggregates, this value was 5.4% and 5.7%, for the TNFɑ-treated and control samples, respectively. For the pre-synaptic aggregates, this value was 4.7% for the TNFɑ treatment and 3.2% for the vehicle treated controls, meanwhile ∼5.9% of the post-synaptic aggregates were fibril-like in the TNFɑ-treated iNeurons and this value was 5.7% for the controls (**Figure 2d**).

We then continued our investigation by characterising the AT8+ tau aggregates. Their total number did not differ significantly between the treatment conditions in terms of aggregate quantity. This was mostly driven by the fact that over 90% of the AT8+ tau aggregates were found in the extra-synaptic compartments, which did not differ between the treatments (**Figure 2e**), also explaining the lack of a significant difference between the vehicle and TNFɑ-treated neuronal lysate measures in SIMOA. However, when we quantified the pre- and post-synaptic aggregates specifically, there was a significant difference in the pre-synapse, with more aggregates in the TNFɑ-treated iNeurons, yet this difference was not present in the post-synapse (**Figure 2e**). These results show the importance of studying the spatial distribution of the aggregates inside the neurons, as little change occurs in the total number of aggregates between the treatment conditions, whereas significant changes occur in the pre-synapse. Since the presence of aggregates is not disease specific and they are also observed in control cases^27,30,38,50^, these results suggest that the pre-synapse is more vulnerable to pathological aggregation due to chronic inflammatory stress.

The average length of the extra-synaptic AT8+ aggregates was ∼255 nm and did not differ between the treatment conditions, meanwhile, the length of the pre-synaptic aggregates was increased with TNFɑ treatment from an average of ∼210 nm to ∼320 nm. On the other hand, there was a trend of shorter aggregates in the post synapses in the TNFɑ-treated iNeurons (average of ∼250 nm in the control cases vs ∼200 nm in TNFɑ; **Figure 2f**). Eccentricity of the extra-synaptic aggregates increased slightly with the TNFɑ treatment, meanwhile the effect of treatment was greater on the pre-synaptic aggregates, but no difference was observed in the post-synapse (**Figure 2g**). Similar to MC1, we once again calculated the proportion of fibrillar AT8+ aggregates, using the same parameters. For the extra-synaptic aggregates, this value was ∼8.0% in the vehicle treated neurons and ∼10.3% for TNFɑ treatment. Meanwhile, for the pre-synaptic aggregates, this value was only ∼1.0% in the control neurons and increased to ∼13.2% in the TNFɑ treated cells. Lastly, in the post-synapse, the change was much smaller, with a value of ∼4.8% in the vehicle treated controls and ∼5.7% in the TNFɑ treated cells (**Figure 2h**).

Following AT8, we characterised the PHF13.6+ tau aggregates, which are considered to be at later stages of aggregation. Similar to the other antibodies, the total number of aggregates did not differ between the treatment conditions. Further investigation of different compartments confirmed this as the extra-, pre-, and post- synaptic regions also did not show any significant difference (**Figure 2i**). Similar to their quantity, the length of the PHF13.6+ tau aggregates also did not differ between the TNFɑ-treated and control iNeurons in the extra-, pre-synaptic compartments, but the PHF13.6+ tau aggregates were longer in the post-synapses of the TNFɑ-treated iNeurons (**Figure 2j**). Lastly, the eccentricity of the PHF13.6+ aggregates also did not differ between the extra-, pre-, and post-synaptic compartments (**Figure 2k**). The proportion of fibrillar aggregates was ∼7.0% and 7.8%, 6.9% and 4.3%, 5.3% and 5.6% in the TNFɑ and vehicle treated samples for the extra-synaptic, pre-synaptic, and post-synaptic compartments, respectively (**Figure 2l**).

Collectively, these results suggest that when challenged with an inflammatory cytokine, AT8-positive tau aggregation increases specifically in the pre-synaptic compartments, and the aggregates formed largely remain non-fibrillar. Meanwhile, no difference between the treatments is detected with the early confirmation change specific antibody MC1 or the late phosphorylation specific antibody PHF13.6, but the tau aggregation patterns agree with the specificity of these antibodies, since the greatest number of aggregates are detected with MC1 (**Figure 2a**), followed by AT8 (**Figure 2e**), and the least number of aggregates are detected with PHF13.6 (**Figure 2i**), suggesting that these aggregates that are related to different disease stages are forming in the iNeurons in a temporal order that resembles the AD brain.

Combining STED microscopy with pre- and post-synaptic markers also allowed us to count the number of pre-and post-synaptic terminals in the TNFɑ- and vehicle-treated iNeurons to determine synaptic loss, along with quantifying the size of the synaptic terminals in terms of diameter and area. While the pre-synaptic count was significantly higher in the TNFɑ treated samples, there was no difference in the post-synaptic count (**Figure 3a**). Meanwhile, the diameter (**Figure 3b**) and area (**Figure 3c**) of the synaptic terminals were reduced in the TNFɑ-treated samples, especially in the pre-synapse. Together, these results suggest that while there is no synaptic loss in the TNFɑ-treated iNeurons, there is a reduction in the size of the synapses, which may be compensated by making more synaptic connections.

**Figure 3.**
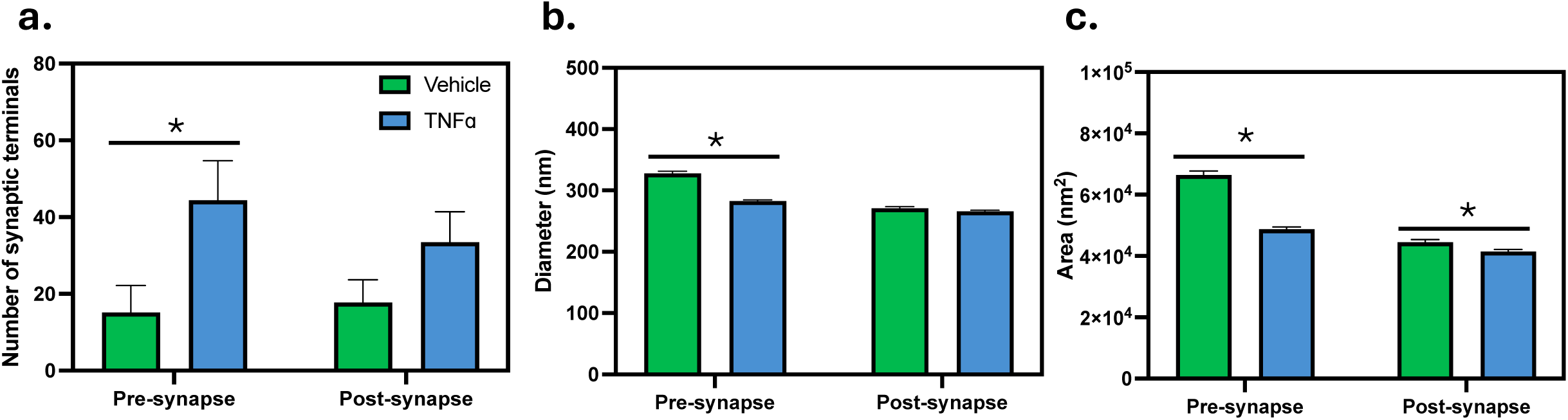
Synaptic terminals characterised by STED microscopy. Number (a), diameter (b), and area (c) of pre- and post-synaptic terminals of iNeurons quantified using pre-synaptic marker Synapsin1/2 and the post-synaptic marker Homer1.

Given that the primary finding of STED imaging was a greater number of longer and more fibril-like AT8+ tau aggregates in the pre-synaptic terminals of the TNFɑ-treated neurons compared to their post-synaptic compartments, we validated this finding in human post-mortem AD brain samples. Using orbitofrontal cortex samples from Braak stage 3 brains that we have characterised previously^38^, we imaged the AT8+ tau aggregates in the pre- and post-synaptic compartments using the same methods we have used on the iNeurons. This disease stage was selected since synaptic pathology is observed in this region without tangle pathology, thus representing an early disease stage. Agreeing with the iNeuron findings, there was a greater number of aggregates in the pre-synaptic compartments compared to the post-synapse (**Figure 2m**). While the aggregate length in the brain (235 nm) was shorter than in the iNeurons (258 nm), the aggregates in the pre-synapses of AD brains (266 nm) were longer than the post- and extra-synaptic aggregate (∼230 nm) similar to the findings from iNeurons (**Figure 2n**). Lastly, we calculated the percentage of the fibril-like aggregates in the AD brain samples using 250 nm as the length cut-off and 0.9 as the eccentricity cut-off, which showed that ∼5.9% of the pre-synaptic aggregates are fibril-like, while none of the aggregates within the post-synapse met these criteria (**Figure 2o**).

### 2.4 SynPull

Characterisation of the tau aggregates in the TNFɑ- and vehicle-treated iNeurons with STED microscopy showed that the number of AT8+ tau aggregates increases in the pre-synaptic compartments under an immune challenge. To further study this, we used SynPull, which is a method we have developed that allows the characterisation of pre-synaptic aggregates with a resolution limit close to 30 nm. This improved resolution over STED enables more accurate morphological characterisation of aggregates, along with their co-localisation in the same synaptosome with markers associated with synaptic loss (phosphatidylserine), synaptic protection (CD47), and synaptic dysfunction (synaptogyrin), which we have also previously studied in human AD brain samples^38^.

Agreeing with the STED results, the number of AT8+ tau aggregates were increased in the synaptosomes harvested from the TNFɑ-treated iNeurons (**Figure 4a**), along with a trend of increased fraction of synaptosomes with one or more aggregate (**Figure 4b**). The AT8+ tau aggregates found in the TNFɑ-treated samples were also longer (**Figure 4c**), but their area (∼3800 nm^2^) and eccentricity (**∼**0.75) did not increase significantly with treatment.

**Figure 4.**
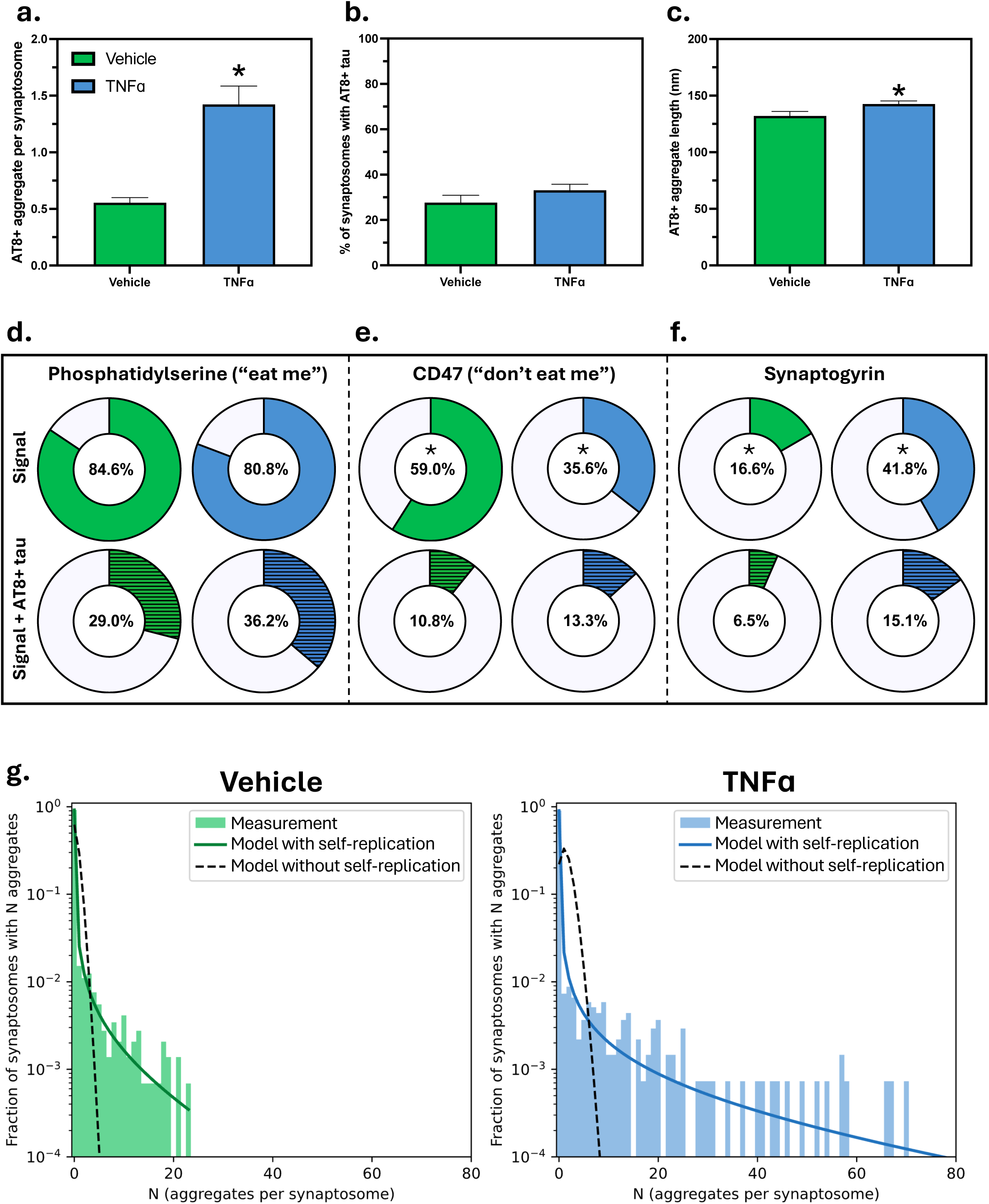
SynPull measurements of the aggregates. Number of AT8+ tau aggregates per synaptosome (a), fraction of synaptosomes with one or more AT8+ tau aggregate (b), and AT8+ tau aggregate length (c) measured by SynPull. Fraction of synaptosomes with exposed “eat me” signal phosphatidylserine (d), don’t eat me signal CD47 (e), and synaptic dysfunction marker synaptogyrin (f), and their co-localisation with AT8+ tau aggregates. The mathematical model provides insight for the mechanisms of aggregate formation (g): the bars show the experimentally measured number of aggregates per synaptosome. The dashed line shows the misfit of a model that does not include self-replication (Poisson distribution). The solid line shows the maximum likelihood fits of the model including self-replication calculated using Equation 1.

Around 82% of the synaptosomes were positive for exposed phosphatidylserine (“eat me” signal) and in ∼32% of the synaptosomes exposed phosphatidylserine signal co-existed with at least one AT8+ tau aggregate, yet neither value differed between the treatment conditions (**Figure 4d**). On the other hand, the fraction of synaptosomes with the “don’t eat me” signal differed significantly between TNFɑ-treated (∼35.6%) and control samples (∼59.0%), while the co-localisation of this signal with AT8+ tau was ∼11% in both conditions (**Figure 4e**). Lastly, the fraction of synaptosomes with synaptogyrin was increased in TNFɑ-treated samples (∼41.8%) compared to the controls (∼16.6%), along with the co-localisation of this signal with AT8+ tau aggregates (TNFɑ: ∼15.1%, vehicle: ∼6.5%; **Figure 4f**).

In addition to the aggregate morphology, SynPull also allows us to determine the distribution of the number of aggregates per synaptosome, which in turn can give insight into the mechanisms that produce these aggregates. We have previously shown that there is evidence of aggregate replication in synaptosomes from AD brains, i.e. existing aggregates trigger the formation of more aggregates^44^. Here we explicitly consider the stochastic effects that dominate in small volumes, such as a synapse, to derive the resulting distribution of aggregates per synaptosome for a simple model of aggregate formation, replication, and removal. The model includes three processes: the self-replication of aggregates, the removal of aggregates, and a process that accounts for both primary nucleation (formation of aggregates without the involvement of existing aggregates) and the import of aggregates into the synaptosome from extra-synaptic space. We refer to the latter process as spontaneous appearance (details are given below in Methods and supplemental material). This is the simplest model for aggregate self-replication, and it predicts that the aggregate number per synaptosome follows a negative binomial distribution. By contrast, a model that does not include self-replication, i.e. just spontaneous appearance and potentially removal, predicts a Poisson distribution. **Figure 4g** shows that a Poisson distribution is inconsistent with the experimentally observed distribution, whereas a negative binomial distribution fits the data well.

These fits also allow us to extract the relative rates of the different processes and compare them between the iNeurons treated with TNFɑ and controls. We found that the rate of self-replication was about a factor of 40 higher than the rate of spontaneous appearance, both in untreated cells (33x higher) and in cells treated with TNFɑ (40x higher). Because the measurements were performed at a single time-point, absolute rates are difficult to obtain, and we instead focus on ratios of rates, which are more robust. However, it can be informative to obtain order of magnitude estimates: for this we assume that the aggregation process took place over the course of approximately 20 days (the age of the cells). Based on this assumption, we can estimate the doubling time due to replication to be four days or less (see supplemental material for details). By contrast, the doubling time obtained from the total accumulation of aggregates in the brain are on the order of years^51^. However, these whole brain rates include both the intra-cellular replication we measure here and the transfer of pathology from one cell to another. Moreover, neurons in brain are surrounded by glia, which can be both supportive and detrimental and thus show different aggregation patterns than the iNeuron cultures that lack glial cells. A more comparable experiment can be found in Dimou *et al*^52^., where we monitored the accumulation of aggregates over time and found doubling times of between 1 and 10 days, depending on the cell type and conditions. The similar orders of magnitude in these two systems, despite the very different methods of measurement, is suggesting that the results are biologically valid.

## 3. Discussion

Pathophysiology of AD may begin decades before the behavioural symptoms appear^53^, making it necessary to study the early changes. However, this is a difficult task due to the lack of suitable models of sporadic AD. One of the primary early drivers of AD is inflammation: it has been shown that chronic inflammation can initially trigger and then worsen beta-amyloid and tau aggregation^54^, while these aggregates can in return induce microglia-driven inflammation^55,56^ creating a pathological positive feedback loop. Indeed, long-term non-steroidal anti-inflammatory drug use has been linked to reduced risk of developing AD^57^. TNFɑ is a pro-inflammatory cytokine, primarily released from microglia in the brain^58^ as a response to cellular stress and damage to initiate a healing process, thus beneficial in the short-term. However, if the release of TNFɑ becomes chronic, the pathological inflammation can lead to neuronal dysfunction and loss^59^.

Here, we used an accelerated human-derived cell-culture model, which shows neuronal and synaptic markers within 10 days *in-vitro*^42^. Treating these iNeurons with TNFɑ for 18 days and then characterising the nanoscopic tau aggregates formed by them led to three main conclusions: (1) chronic inflammatory stress leads to increased AT8+ tau aggregate accumulation, and this effect of TNFɑ is limited to the pre-synapses. The pre-synaptic terminals also shrank, suggesting functional deficits^60–62^. (2) As revealed by our mechanistic models, the increased aggregation in the pre-synapses is driven both by new aggregate formation and replication, however the dominant mechanism is replication, with an approximate aggregate doubling time of under four days. (3) In terms of their size and shape, the pre-synaptic aggregates differ from the post-synaptic and extra-synaptic aggregates (which were more similar to each other), suggesting a lack of synaptic aggregate transfer in this system.

One key question is why the pre-synapse is most susceptible to pathological aggregation, although TNF receptors are expressed in both pre- and post-synapses, as well as the soma^63^. Our analysis shows that the increased numbers of aggregates at the pre-synapse are formed predominantly by replication rather than nucleation. Application of TNFɑ leads to the formation of more aggregates. One explanation for this increase in apparent rate of formation could be an increase in the pre-synaptic concentration of phosphorylated tau in the presence of TNFɑ. As a microtubule associated protein concentrated on the axons, tau can translocate to the pre-synaptic terminals under pathological conditions and bind to neurotransmitter vesicles, contributing to the synaptic pathology^64,65^. Similarly, reduced removal rates at the synapse could also lead to an increase in the rate of aggregate accumulation. While autophagosomes are often formed in the pre-synaptic terminals, they need to be carried back to the soma where the lysosomes are concentrated, with retrograde transport^66^. This process is disrupted in AD and under chronic inflammation^67,68^, leading to protein degradation dysfunction in the pre-synapse and accumulation of toxic aggregates. Both scenarios are also consistent with our previous analysis of the length distribution of tau aggregates in synaptosomes of AD patients, which showed that there was increased formation or reduced removal at synapses compared to the rest of the neuron, explaining why aggregation may occur preferentially at synapses^38^. Moreover, due to its high energy demand, the pre-synapse is rich in mitochondria and vulnerable to oxidative damage -more than other parts of the neuron^69–71^.

Alongside increased tau aggregation, other pathological outcomes of chronic inflammatory stress are the reduction of pre-synaptic size and synaptic protective signal CD47^72,73^, and increased synaptogyrin, which has been linked to pre-synaptic dysfunction^74^, collectively suggesting pre-synaptic pathology. While our data does not provide conclusive causative links between this pathology and tau aggregation in either direction, it will be highly beneficial to investigate these mechanisms in future studies, which can then be therapeutically targeted. Interestingly, exposed phosphatidylserine levels did not differ between TNFɑ-treated and control samples, as the levels were overall quite high (80% of the synaptosomes), suggesting that the iNeurons are stressed, potentially due to the lack of supportive glia in the cultures^75–77^, which is also likely the reason why we did not observe synaptic loss, which is mostly mediated by microglia and astrocytes^41^.

While tau fibrils have been demonstrated to seed aggregation leading to an increase in the amount of aggregate present in a sample^78^, the number of studies that have directly characterised the aggregates formed after seeding and shown that they directly increase in number in a cell is limited^79^ and has not focused on oligomeric aggregates. While pre-synaptic tau oligomer accumulation was previously demonstrated by our group^38,44^ and others^80^, to the best of our knowledge, replication of tau aggregates that have formed spontaneously, rather than been introduced as seeds, has not been observed at synapses, so our results are providing the first direct evidence that this is possible. Importantly, it means that once an aggregate forms at a synapse the aggregate number can rapidly increase.

Colom-Cadena et al.^80^ and McGeachan et al.,^81^ reported oligomeric tau in both the pre-and post-synaptic terminals from AD and progressive supranuclear palsy, respectively, with elegant studies using array tomography. The observation of tau in pre- and post-synaptic pairs with a frequency higher than chance (i.e. higher frequency of seeing tau aggregates in the post-synapse if the pairing pre-synapse also contained aggregates) was interpreted as evidence for synaptic spread of oligomeric tau, yet the pre- and post-synaptic aggregates were not morphologically characterised in these studies. For a number of reasons our results here - obtained using methods with higher sensitivity, so we can detect more synapses with small aggregates, combined with the capability to gather morphological information from individual aggregates, indicate that synaptic spread is at least not the dominant mechanism in the iNeurons at the stage we studied them and the human AD pre-frontal cortex. First, the increased aggregation is only seen in the pre-synapse and no other part of the iNeurons, including the post-synapse. Importantly, in terms of their size and shape, the post-synaptic AT8+ tau aggregates differ from the pre-synaptic ones as they are shorter and less fibril-like, both in the iNeurons and AD brain. Therefore, even though tau aggregates exist on both sides of the synapse, they seem to be morphologically different species. While cannot exclude the possibility that the pre-synaptic aggregates may be taken up in the post synapse and then modified, our data provides stronger evidence for early pathological aggregation of AT8+ tau in the pre-synapse, primarily driven by local replication, rather than synaptic spread. Indeed, aggregate-specific SIMOA showed an increase of AT8+ tau aggregate levels in the conditioned media from the TNFɑ-treated iNeurons, which was not accompanied by intra-cellular aggregate levels, together suggesting that the aggregates made by the treated iNeurons can be released by the pre-synaptic neuron but not reuptaken by the post-synaptic neuron. Instead, they may activate microglia^82^, which can then increase tau phosphorylation in neighbouring neurons through NLRP3 inflammasome^83^, which also explains the higher rates of synaptic tau observed in neurons with in proximity.

Despite these novel biological insights and methodological strengths, our work does have a number of limitations, which needs to be addressed in future studies. First of all, cultured neuronal systems, including the iNeurons used here, often only express the developmentally dominant 3R isoforms of tau, lacking the 4R isoforms, unlike the adult human brain^84^. Our methods are limited in spatial orientation, and synaptic pairs cannot be studied. Moreover, we have only used a single cytokine in this study (TNFɑ), due to its heavy involvement in AD pathogenesis and high levels of detection in the blood and cerebrospinal fluid of early AD and mild-cognitive impairment cases^85,86^, yet other cytokines such as interleukin 1 beta and interferon gamma are also involved in the pathophysiology of AD^87,88^. Lastly, our cultures only contained neurons, meanwhile other cell types such as astrocytes and microglia are also heavily involved in AD. The lack of astrocytes in our samples may lead to reduced clearance of glutamate from the synapse, leading to excitotoxicity and stress in the control cells^89^, which was detected by our data modelling and indicated by high levels of exposed phosphatidylserine. Moreover, it has been suggested that astrocytes are necessary for the microglia-induced synaptic elimination in AD^90^. Since our system lacks both of these types of glial cells, elimination of synapses due to tau pathology cannot be studied, which explains the lack of synaptic loss, even in the presence of pathological alterations in synaptic morphology. As such, future work including multiple cells types co-cultured will be useful to identify the downstream mechanisms of synaptic tau aggregation.

In conclusion, our results are providing strong evidence for pre-synaptic oligomeric tau aggregation being an early pathological event in neurons driven by chronic inflammation. While the appearance of a pre-synaptic aggregate remains a rare event, once a synaptic aggregate forms, replication dominates over nucleation and influx for the formation of new aggregates. The morphological differences between pre-synaptic aggregates and ones from other parts of the neuron suggests that this is a local pathology. While the patho/physiological outcomes of tau aggregation need to be further studied in future experiments, these results provide a mechanism of how tau aggregation occurs in neurons and causes synaptic dysfunction and highlights the importance of inflammation in driving this process.

## 4. Methods

### 4.1 Maintenance of induced pluripotent stem cells

Three independent induced pluripotent stem cell (iPSC)-derived neurons were used in the experiments. (1) Commercially available ioGlutamatergic Neurons (Bit Bio Ltd., Cat. io1001), (2) human Kolf2 iPSCs originally generated by the HipSci (Human Induced Pluripotent Stem Cells) initiative and available from Public Health England/ECACC (HPSI0114i-kolf_2, Cat. 77650100), and (3) i3 neurons^91^ (from Michael E. Ward, National Institute of Health), which are hiPSCs with stable integration of NGN2 into a safe-harbour locus under a doxycycline-inducible promoter^91^. Differentiation and growth protocols of these lines are detailed below.

For the iPS cell lines (2) and (3), human iPSCs were cultured in StemFlex media (Thermo Fisher Scientific, Cat. A3349401) containing 1% penicillin-streptomycin (Fisher Scientific, Cat. 15-140-122) and kept in a humidified incubator at 37℃ and 5% CO_2_. A complete media change was performed every second day, and cells were passaged every 5-7 days, depending on confluency. For cell passaging, the medium was aspirated, cells were rinsed once with Dulbecco’s phosphate-buffered saline (DPBS) without calcium and magnesium (Life Technologies, Cat. 14190-094) and incubated in ReLeSR (Stem Cell Technologies, Cat. 100-0483) at room temperature for 1 minute. ReLeSR was then aspirated, and cells were incubated in a thin layer of remaining liquid for a further 4 minutes. Cell detachment was facilitated by forceful washing with DMEM/F12 (Life Technologies, Cat. 31330-038). The resultant cell aggregates were transferred into a 15 mL falcon tube containing 5 mL of DMEM/F12 and were left to precipitate. Approximately 100 μL of cell suspension was plated onto tissue-culture treated 6-well plates (Corning) pre-coated with Geltrex (Fisher Scientific, Cat. A1413302), to achieve a split ratio of ∼1:10. For the coating of plates, 10 μL of Geltrex was diluted in 1 mL of ice-cold DMEM/F12 per well of a 6-well plate. Plates were incubated at 37°C for at least one hour. Immediately before plating, Geltrex solution was aspirated and replaced with fresh pre-warmed cell-containing StemFlex medium.

### 4.2 ioGlutamatergic Neurons

The parental iPSC line with normal karyotype used by the manufacturer originated from skin fibroblasts obtained from an adult Caucasian male donor (55–60 years of age). These neurons were cultured following the manufacturer’s deterministic cell programming protocol using opti-ox™. During Phase 0 (Induction) -which was performed entirely at the manufacturer’s facility, iPSCs underwent a transcription-factor-driven programming protocol that irreversibly exits pluripotency and commits cells to a glutamatergic neuronal fate before cryopreservation. Upon receipt of cryopreserved cells, neurons were thawed according to the supplier’s protocol and transitioned into Phase 1 (Stabilisation). Cells were resuspended in complete glutamatergic neuron medium supplemented with 1 μg/mL doxycycline (comp:GN+D) and plated onto poly-D-lysine (PDL) and Geltrex-coated vessels at a density of ∼30,000 cells/cm². Over the first 96 hours, cultures were maintained in doxycycline-containing medium, between 48–96 hours to reinforce neuronal induction. Media handling was performed gently using micropipettes to prevent stress-induced detachment. From Phase 2 (Maintenance; day 4 onwards), cells were transferred to doxycycline-free complete neuronal medium (comp:GN) and maintained with half-volume medium changes every 48 hours. Under these conditions, ioGlutamatergic Neurons exhibited stable attachment, healthy morphology, and sustained maturation.

### 4.3 Human Kolf2 iPSCs

Originally generated by the HipSci (Human Induced Pluripotent Stem Cells) initiative and available from Public Health England/ECACC (HPSI0114i-kolf_2, Cat. 77650100), Kolf2 cells engineered to carry the NGN2 opti-ox cassette were differentiated into excitatory glutamatergic neurons following established forward-programming protocols^42,92^. In brief, ∼70% confluent iPSCs were dissociated into single cells using Accutase (ThermoFisher, Cat. A1110501) and replated onto Geltrex-coated plates in StemFlex medium supplemented with ROCK inhibitor (Tocris, Cat. 1254/10). After 24 hours, on DIV 0, neuronal forward programming was initiated by replacing the medium with DMEM/F-12 supplemented with Glutamax (100x, ThermoFisher Scientific, Cat. 35050038), Non-Essential Amino Acids (100x, Life Technologies Ltd, Cat. 11140-050), 50 μM 2-Mercaptoethanol (Thermo Scientific, Cat. 31350010), 1% Penicillin/Streptomycin and 1 μg/mL Doxycycline (Stem Cell Technologies, Cat. 72742). At DIV3, cells were again dissociated with Accutase and replated at assay-dependent densities onto Poly-D-Lysine (PDL)/Geltrex-coated plates in Neurobasal (ThermoFisher, Cat. 21103049) medium supplemented with Glutamax (100x), B27 (50x), 10 ng/mL BDNF (PeproTech, Cat. PHC7074), 10 ng/mL NT3 (R&D Systems, Cat. 267-N3), 1% Penicillin/Streptomycin, and 1 μg/mL Dox. Cultures were maintained with daily medium changes until DIV7, followed by 50% medium changes every other day until readout. Dox was withdrawn at DIV6 post induction.

### 4.4 i3 neurons

i3 neurons were cultured in STEM FLEX medium (STEMCELL technologies) on Vitronectin (ThermoFisher Gibco, Cat. A14700)-coated plates (1:100 dilution, 1 hour) and treated with 10µM ROCK Inhibitor (BD biosciences, Cat. 562822) in the first 24 hours after seeding. To induce neuronal differentiation, cells were transferred to Geltrex (ThermoFisher Gibco, Cat. A1413302)-coated plates (1:100 dilution, 1 hour) and kept in STEM FLEX medium supplemented with ROCK Inhibitor for 24 hours. Then, the STEM FLEX was replaced with DMEM/F12 (Gibco, Cat. 11559726) supplemented with 10 µl/ml N-2 supplement (Gibco, Cat. 17502048), 2mM L-glutamine (Gibco, Cat. 25030024), 10 µl/ml non-essential amino acids (Gibco, Cat. 11140035), 50 µM 2-mercaptoethanol, 10 µl/ml Pen-Strep (Gibco, Cat. 15140122) and 1 µg/ml doxycycline (Fischer Scientific, Cat. BP2653-1). This medium was changed daily for two days. On day 3 after induction, the medium was replaced with the maintenance medium Neurobasal (Gibco, Cat. 21103049) supplemented with 20 µl/ml B-27 (Gibco, Cat. 17504044), 2 mM L-glutamine (Gibco, Cat. 25030024), 50 µM 2-mercaptoethanol (Gibco, Cat. 31350010), 10 µl/ml Pen-Strep, 1ug/ml doxycycline, 10 ng/ml NT-3 (Gibco, Cat. 450-03-10) and 10 ng/ml brain-derived neurotrophic factor (Gibco, Cat. 10477253). At day 4, cells were transferred to different kinds of plates, depending on the experiment modality, coated with poly-L-lysine (Bio-Techne, Cat. 3438-100-01) overnight and Geltrex at 1:100 dilution for 1 hour. The media was supplemented with 3 µM uridine (Sigma-Aldrich, Cat. U3003-5G), 1 µM 5-Fluoro-2′-deoxyuridine (Sigma-Aldrich, Cat. F0503-100MG), and 10 µM ROCK inhibitor for 24 hours after seeding. The maintenance medium was subsequently changed every day for 4 days and every other day after DIV 8.

### 4.5 TNFα treatment

To model chronic inflammatory stress in a manner comparable across all differentiation platforms, iPSC-derived neurons were exposed to human recombinant TNFα (PeproTech, Cat. 300-01A-10UG) for a uniform 18-day treatment window. Although the three systems differ in their early induction stages and DIV definitions, as explained below, the TNFα exposure paradigm was harmonised to ensure equivalent total time in culture from the iPSC state.

For ioGlutamatergic Neurons, the initial neural induction (Phase 0) was performed in-house by the manufacturer before cryopreservation. Following thawing in our laboratory (defined as DIV0), cultures were allowed to stabilise for 24 hours, and TNFα treatment commenced at DIV2. Cells were treated for 18 consecutive days with TNFα, with the final experimental endpoint at DIV20. For Kolf2 and i3 neurons, TNFα treatment was initiated at DIV5 (following NGN2 induction in the Kolf2 line) and continued for 18 days until the experimental endpoint at DIV23.

Across all three iPSC-derived neurons, TNFα was applied at a constant concentration of 100 ng/mL, replenished at each medium change to maintain stable TNFα exposure throughout the treatment duration. Phosphate-buffered saline (PBS) was used as the vehicle control. The experimental design ensured that all neuronal cultures experienced the same 18-day TNFα exposure, enabling direct comparison across differentiation protocols with downstream assays.

### 4.6 Cell viability assay

Viability of iPSC-derived neurons were assessed using the RealTime-Glo™ MT Cell Viability Assay (G9711, Cat. Promega) according to the manufacturer’s instructions. Neurons were seeded at 10 × 10³ cells per well into poly-D-lysine (PDL)/Geltrex-coated 96-well plates and maintained under standard culture conditions until assay endpoint. At the time of assay, RealTime-Glo reagent was freshly prepared in neuronal culture medium, added directly to the wells, and luminescence was measured at defined intervals using a Tecan Infinite 200 Pro microplate reader, reflecting cellular metabolic activity. All readings were background-corrected against medium-only controls and compared across conditions.

### 4.7 Conditioned media and cell lysate preparation and SIMOA analysis

For cell lysates and conditioned media preparations, approximately 200 × 10³ cells were seeded per well of a 6-well plate. For all neuronal lines (ioGlutamatergic Neurons, Kolf2 NGN2 iNeurons, and i3 neurons), conditioned media were collected at experimental endpoints into protein LoBind tubes (Eppendorf, Cat. 022431064) to minimise protein adsorption. Samples were centrifuged at 300 × g for 5 min at 4 °C to remove cell debris, and the clear supernatant was transferred into fresh LoBind tubes, snap-frozen on dry ice, and stored at -80 °C until analysis. For preparation of cell lysates, cultures were washed once with ice-cold PBS and lysed in buffer consisting of PBS with 1% Triton X-100 supplemented with protease (Roche, Cat. 11873580001) and phosphatase inhibitors (Roche, Cat. 4906845001). Lysates were incubated on ice for 30 min, followed by centrifugation at 14,000 × g for 10 min at 4°C^37^. The resulting supernatants were transferred to LoBind tubes and stored at -80°C until SIMOA analysis.

AT8-positive tau aggregate-specific single-molecule array (SIMOA) analysis of the media and cell lysate samples were performed as previously described^26^. In brief, the samples were incubated with SiMoA^®^ Homebrew carboxylated beads (Quanterix; Cat.104006) functionalised with AT8 antibody on plate shaker at 30°C at 800 rpm for 30 minutes. The plate was then washed by the SiMoA washer with the provided buffers. Then, the aggregate bound beads were incubated with biotinylated detection antibody AT8 for 10 minutes on the plate shaker at 30°C with 800 rpm. Following another wash, the beads were incubated with 50 pM streptavidin-beta galactosidase (SBG) for 10 minutes at 30°C at 800 rpm. Finally, the plate was washed by the SiMoA washer and analysed on the SR-X™ Biomarker Detection System.

### 4.8 Immunofluorescence staining and STED microscopy

Neurons derived from ioGlutamatergic Neurons, Kolf2 NGN2 iNeurons, and i3 Neuron differentiation platforms were processed for immunocytochemistry following a previously described protocol^29,93–96^. Briefly, cells were seeded at a density of 60 × 10³ cells per well on poly-D-lysine (PDL)/Geltrex-coated #1.5H coverslips (Paul Marienfeld GmbH, Cat. 0117530)) in 24-well plates and maintained throughout differentiation until the downstream assay endpoint. At the endpoint, samples were fixed with 4% paraformaldehyde and 4% sucrose in PBS, then permeabilised with 0.3% Triton X-100 in PBS for 10–15 minutes. They were subsequently incubated in a blocking solution containing 10% normal goat serum (Abcam, Cat. ab7481) and 0.3% Triton X-100 in PBS for 30–45 minutes to minimise non-specific antibody binding. Cells were then incubated for one hour at room temperature with combination of primary antibodies anti-MAP2 (1 ng/μL; Abcam, Cat. ab5392), anti-Synapsin1/2 (5 ng/μL; Synaptic Systems, Cat. 106 004), Homer1 (5 ng/μL; Synaptic Systems, Cat. 160 003), anti-Phospho-Tau (Ser202, Thr205), AT8 (1 ng/µL; Invitrogen, Cat. MN1020), anti-Phospho-Tau (Ser396), PHF13.6 (2.5 ng/µL; Invitrogen, Cat. 35-5300), Phospho-Tau (Thr231), AT180 (1 ng/µL; Invitrogen, Cat MN1040), and MC1 (1:100, Antibody Systems, Cat. RHC82420) diluted in primary antibody solution containing 3% normal goat serum and 0.3% Triton X-100 in PBS. After three washes in PBS, appropriate fluorophore-conjugated secondary antibodies, Abberior STAR RED (AbberiorGmbH, Cat. STRED-1001-500UG), Alexa Fluor™ 594 (Life Technologies Limited, Cat. A11076 and A11037), Alexa Fluor™ 488 (Life Technologies Limited, Cat. A78948) (all 1:500 dilutions) in secondary antibody solution containing 3% normal goat serum and 0.3% Triton X-100 in PBS were applied for one hour at room temperature and subsequently washed with PBS three times. Coverslips were mounted onto glass slides using an antifade mounting medium, ProLong™ Diamond Antifade Mountant (Invitrogen, Cat. P36961). Samples were stored in the dark until further imaging.

We used a commercial stimulated emission depletion (STED) inverted microscope (Abberior STED Expert Line Super Resolution Microscope, Abberior Instruments GmbH, Göttingen, Germany). This laser scanning microscope provides multiple super-resolution and confocal channels. It is based on a fully automated Olympus IX83 microscope platform with a 100x (1.4 NA) oil immersion objective (UPLSAPO 100XO) and a high-precision Ultrasonic Stage (IX3-SSU). STED and confocal image acquisition was performed with pulsed 488 nm, 561 nm and 640 nm laser excitation (1mW@40MHz, 250µW@40MHz & 1mW@40MHz maximum output power respectively) and depletion at 775nm (3W @40MHz maximum output power). The fluorescence was collected with Avalanche Photodiode Detectors (APDs) operating in single photon counting mode. Images were taken for a region of interest (750 pixels x 750 pixels) with a pixel size of 20nm. The pinhole was set to 1.0 AU.

For dual-colour STED imaging, the same depletion laser was used in lateral depletion mode while the excitation wavelength was adapted to the fluorophore. The STED was performed using a pulsed depletion laser at 775 nm wavelength with a gating on delay of 750 ps and a dwell time of 30 μs. The APD1 detector (detection 650 nm to 763 nm, excitation 640 nm) was used for Abberior Star Red, APD2 detector (detection 584 nm to 630 nm, excitation 561nm) was used for Alexa Fluor 594, and APD3 detector (detection 498 nm to 551 nm, excitation 488 nm) was used for Alexa Fluor 488. The laser powers were adjusted to 25% and 8% of their respective total power for 561 nm and 640 nm, respectively. A depletion laser power of 8% and 7% of the total power of 775 nm lasers was used for depleting 561 nm and 640 nm, respectively. The laser power for the Alexa 488 was adjusted to 5 % of the total power. The Alexa Flour 488 images were recorded only in the confocal mode. The image acquisition was controlled by the Imspector software. The software is proprietary and designed by Abberior Instruments for the control and gathering of measurement data. A comparable approach was employed in earlier work^93,95,97^. Before commencing imaging sessions for the experimental categories, precise alignment of the STED doughnut with the excitation beam was conducted as an essential preliminary step. We employed the Abberior STED Expert Line Super Resolution Microscope, Abberior Instruments, equipped with programmable adaptive optics elements such as Spatial Light Modulators (SLM). The beam shape for the two-dimensional doughnut was created using SLM. Abberior’s fluorescent adjustment sample (four-color TetraSpeck 100-nm microspheres, NP-3011) facilitated standardised alignment. The STED beam position was corrected with respect to the confocal signal by adjusting the grating of the SLM, specifically the grating X and grating Y controls. By tuning the frequency of the grating, the steering of the STED beam was precisely adjusted to optimise the overlay between the STED and excitation beams.

To minimise sampling bias, imaging fields were selected using a randomised approach within MAP2-positive regions, and acquisition was performed blinded to sample identity. We ensured that each comparable group was configured with the same settings. This approach helped minimise any potential subjective bias in regions of interest selection. To detect different phosphorylated variants of tau, we used primary antibodies including AT8, AT180, and PHF13.6, while the conformation-dependent MC1 antibody was used to label aggregated tau. For subsynaptic localisation studies, tau and phospho-tau signals were compared with the distribution of pre- and postsynaptic markers using anti-Synapsin1/2 and anti-Homer1 antibodies, respectively. Fluorophore-conjugated secondary antibodies included Abberior Star Red for Tau-related targets and Alexa Fluor 594 for synaptic markers. Alexa Fluor 488 was used to visualize MAP2.

### 4.9 STED super-resolution cluster analysis and detection of synapses

Image analysis was performed using a custom Python script. Raw STED image files (.msr format) were processed using the “obffile” library, enabling extraction of the relevant STED channels corresponding to pre- or post-synaptic markers (AF594) and tau (STAR Red). STED images of the pre- and post-synaptic markers Synapsin 1/2 and Homer1 were used to define synaptic regions and to distinguish them from extra-synaptic areas.

To isolate, synapse and tau candidates, the image background was first removed using an intensity threshold. A white top-hat filter was then applied to enhance the bright spots against the remaining background. To remove noise, the images underwent morphological opening (erosion followed by dilation) and the removal of small objects with an area of fewer than 5 pixels. This processed image was binarised to create a mask where bright regions (labeled ‘1’) represented the detected structures and dark regions (labeled ‘0’) represented the background. Any remaining internal gaps within the detected spots were filled using a binary fill holes operation to ensure each spot was a solid area. The identified synapses were then filtered by size, and only those with a major axis length between 150 and 800 nm were retained for further analysis. This filtering criterion was chosen based on the fluorescent puncta characteristics appropriate for detecting synaptic structures, as estimated from synaptic dimensions reported by Harris and Stevens^98^, and is described earlier^29,96^. The identified pre- or post-synapses were subsequently used for analysing tau clusters. A similar workflow was applied to the Tau STED images. The “regionprops_table” function from the scikit-image package was used to calculate the centroids of the aggregates. An aggregate was considered colocalised to a pre-synapse or a post-synapse if their centroids overlapped with the identified pre- or post-synapse mask. These were classified as presynaptic or postsynaptic aggregates, respectively while those falling outside this criterion were designated as extra-synaptic aggregates. For each tau aggregate, the following quantitative parameters were computed: area, and major and minor axis lengths using the “regionprops_table” function.

### 4.10 Synaptosome preparation and imaging

iPSC-derived neurons from the three independent lines were washed once with ice-cold PBS, detached from culture plates by gentle scraping, and homogenised on ice in Syn-PER™ Synaptic Protein Extraction Reagent (ThermoFisher Scientific, Cat. 87793). Homogenates were transferred to protein LoBind tubes and centrifuged at 1,200 × g for 10 min at 4°C to pellet cellular debris. The resulting supernatant was centrifuged at 15,000 × g for 20 min at 4°C to isolate the synaptosome fraction. Pellets were gently resuspended in Syn-PER buffer and stored at -80°C until further analyses.

Imaging of the synaptosomes was performed as previously described^44,97^. In brief, synaptosome samples were incubated on the PEG coated SiMPull coverslips^99^ with the anti-NRXN1 capture antibody (10 nm in TBS with 1ug/mL BSA for 15 minutes; abcam, Cat. ab222806) overnight at 4℃. On the following day, the surface-bound synaptosomes were fixed and permeabilised, and then synaptic AT8-positive tau aggregates were stained using Alexa Fluor™ 647 NHS ester labelled anti-AT8 antibody (2 nm in TBS with 1ug/mL BSA; Invitrogen, Cat. MN1020B) along with anti-exposed phosphatidylserine antibody (2 nm in TBS with 1ug/mL BSA; Sigma-Aldrich, Cat. 16-256) for 30 minutes. Lastly, the synaptosome membranes were stained using CellMask™ plasma membrane stain (Thermo Fischer Scientific, Cat. C10045) for 10 minutes (1 : 6,000 by volume in TBS), followed by four rounds of washes with TBS. Direct stochastic optical reconstruction microscopy (*d*STORM) imaging for super-resolution size and shape analyses of the AT8-positive tau aggregates along with diffraction-limited single-molecule imaging for CellMask and exposed phosphatidylserine were performed on a purpose-built total internal reflection fluorescence (TIRF) microscope. Image analysis was performed as previously described^97^.

### 4.11 Model of aggregate number distributions

We consider 3 processes: (1) The self-replication of aggregates, with the rate at which an existing aggregate self-replicates given by κ (please refer to the SI for how this relates to the self-replication rate obtained from more detailed aggregation models^33^). (2) The removal of aggregates, with the rate at which an existing aggregate is removed given by λ. (3) The spontaneous appearance of aggregates, without the involvement of existing aggregates, with rate k. This process accounts for both primary nucleation (formation of aggregates from monomers without the involvement of existing aggregates) and the import of aggregates into the synaptosome from extra-cellular space.

The derivation is detailed in the SI and yields the following expression for the aggregate number distribution:

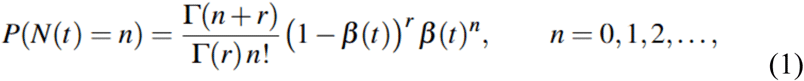

where *n* is the size of the aggregate and the parameters β is given by:

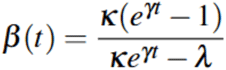

with *γ = κ - λ* and *r = k/ κ*.

The fits of this model were performed using Bayesian inference, i.e. the total probability of a dataset is given by:

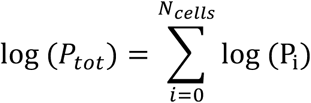

where the sum is over all cells and the individual probability given by the negative binomial distribution above. Priors for the parameters κ, λ and k were assumed to be flat in log space. Errors given on the parameter values are one standard deviation.

## 5. Declarations

## Ethical approval

Human post-mortem brain tissue was acquired from the Cambridge Brain Bank (Cambridge University Hospitals). The Cambridge Brain Bank is supported by the NIHR Cambridge Biomedical Research Centre (NIHR203312). We gratefully acknowledge the participation of all our patient and control volunteers.

## Funding

D.K. receives funds from the UK Dementia Research Institute through UK DRI Ltd, principally funded by the Medical Research Council and holds a Royal Society Professorship. Y.W. is supported by the Early Career Pathway Award (EDDAPA-2024/100006) from the International Alliance for Cancer Early Detection (ACED), Cancer Research UK.

## Competing interests

None to declare.

## Availability of data and materials

Data collected during these experiments will be made available upon reasonable request from the corresponding author. No novel code was generated.

## Author contributions

**S.K.** Conception and design, data collection, image analysis, manuscript preparation. **E.F.** Conception and design, data collection and analysis, manuscript writing. **A.P., V.D., K.B., E.A., and M.R.N.K.** iNeuron development and culturing, sample preparation. **G.N.** Sample preparation. **Y.W.** Image analysis. **J.E.K.** Data collection. **A.Q.** Providing human brain samples, neuropathological characterisation. **G.M.** Data modelling and interpretation, manuscript preparation. **D.K.** Conception and design, data interpretation and manuscript preparation, overall supervision of the project.

## Acknowledgments

We thank Dr. Kristy Halliday from the Cambridge Brain Bank for her assistance on human brain tissue acquisition. STED imaging was performed using the STED microscope funded by BBSRC grant BB/R000395/1 at the Department of Genetics, School of Biological Sciences, with the help of Dr. Martin O. Lenz and Dr. Antonina Kruppa. Deconvolution was performed using the Light Microscopy Core Facility at Cancer Research UK Cambridge Institute. We would also like to thank Ms. Amanda Turner and other members of Bit Bio Ltd. for their support of this project.

## 7. Figure legends

**<u>Supplemental figure 1.</u> Representative images from STED and SynPull.** (a) Nanoscale distribution of AT8-positive AT8+ tau aggregates within functional zones of excitatory post-synapses imaged using STED microscopy. STED image overlays showing the pre-synaptic marker Synapsin1/2 and the post-synaptic marker Homer1 (magenta), together with AT8+ tau (green), in vehicle- and TNFɑ-treated samples. Scale bars: 2.5 µm (top panels) and 0.5 µm (bottom panels). (b) Representative pseudo-colour SynPull images of AT8+ tau aggregates under vehicle- and TNFɑ-treated conditions acquired by *d*STORM microscopy and reconstructed using ACT software^100^. The intensity is pseudo-colour-coded from black (minimum) to white (maximum). Scale bar: 50 nm. Individual aggregates were segmented using MetaMorph software v7.10.5.476 (Molecular Devices)^38,43,44^.

## Notes

**Conflict of interest:** Authors declare no conflict of interest.

### Competing Interest Statement

The authors have declared no competing interest.

