## Supplementary figures and images for "Tau aggregate replication occurs at the pre-synapse of cultured human neurons and increases with application of TNFɑ"

### Supplemental Figure 1

#### Supplemental Figure 1

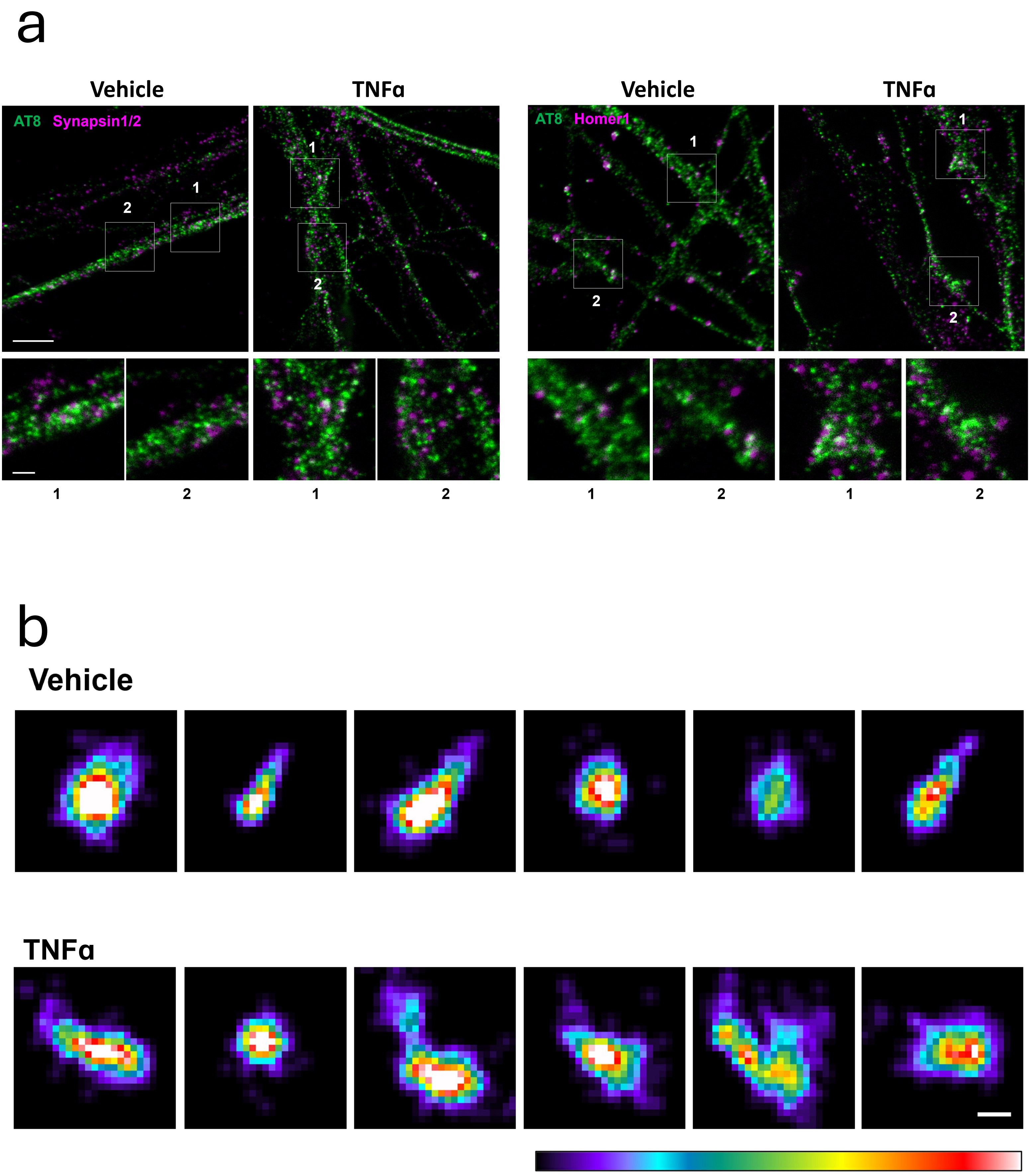
