## Supplemental text for "Tau aggregate replication occurs at the pre-synapse of cultured human neurons and increases with application of TNFɑ"

### Derivation of aggregate number distribution

Consider a population of aggregates in which each aggregate can independently self-replicate at rate $\kappa$ and aggregates are removed at rate $\lambda$. New aggregates enter the system at a constant rate $k$, independently of the current aggregate number, either by spontaneous nucleation or by import. The probability of the system to have $n$ aggregates at time $t$ is denoted by $P_n(t)$. We start from an *empty* system, so the initial condition is $P_0(0) = 1$ and $P_n(0) = 0$ for $n \geq 1$.

Balancing the probability flux gives the master equation

$$\begin{aligned}\frac{dP_0}{dt} &= \lambda P_1 - kP_0,\\ \frac{dP_n}{dt} &= \kappa(n-1)P_{n-1} + \lambda(n+1)P_{n+1}\\ &- \big[(\kappa+\lambda)n + k\big]P_n + k\,P_{n-1}, \qquad n \geq 1. \end{aligned} \tag{1}$$

For readers familiar with our other work on protein aggregation kinetics[1,2,3], note that the states in this master equation denote systems with different aggregate numbers, not the concentrations of different aggregate sizes.

Defining the generating function $G(z,t) = \sum_{n=0}^{\infty} P_n(t)\, z^n$, the initial condition gives $G(z,0) = 1$. Multiplying the master equation by $z^n$ and summing over all $n$ yields

$$\frac{\partial G}{\partial t} = (\kappa z - \lambda)(z-1)\,\frac{\partial G}{\partial z} + k(z-1)\,G, \qquad G(z,0) = 1. \tag{2}$$

This partial differential equation can be solved by the method of characteristics. The characteristic equations for (2) are

$$\frac{dt}{1} = \frac{dz}{-(\kappa z - \lambda)(z-1)} = \frac{dG}{k(z-1)\,G}. \tag{3}$$

We thus obtain

$$G(z,t) = G(z_0,\,0)\cdot \exp\!\Big(\int_0^t k\big(z(s)-1\big)\,ds\Big) = \left[\frac{1-\beta(t)}{1-\beta(t)\,z}\right]^r, \tag{4}$$

where $\gamma \equiv \kappa - \lambda$, $r \equiv k/\kappa$, and

$$\beta(t) = \frac{\kappa(e^{\gamma t}-1)}{\kappa e^{\gamma t} - \lambda}. \tag{5}$$

This is the probability generating function of a negative binomial distribution, NB($r$, $1 - \beta(t)$), for all $t > 0$. Thus

$$P(N(t)=n) = \frac{\Gamma(n+r)}{\Gamma(r)\,n!}\big(1-\beta(t)\big)^r\,\beta(t)^n, \qquad n = 0,1,2,\ldots, \tag{6}$$

### Assumptions and relation to other models of aggregation

We purposefully choose the simplest model here, by simply having a constant self-replication rate and do not consider aggregate growth explicitly (effectively assuming a single aggregate size). The resulting model is simple to understand, matches the measured number distribution well, and parallels early models of cell population dynamics[4]. One could extend this model to explicitly include aggregate growth and multiplication, for example by elongation and secondary nucleation or fragmentation[3]. However, this would increase model complexity significantly, as systems are then characterised not just by the number of aggregates, but by the size of every aggregate in the system, yielding many more states to track. We leave this exercise for future work. However, we note that the aggregating systems can often be approximated well by assuming a single average aggregate size, yielding simpler equations that nonetheless capture the fundamental behaviour well[5,6]. In a model which includes both growth and multiplication, both aggregate number and total aggregate mass tend to grow as $e^{\kappa t}$, where the rate of self-replication $\kappa$ is given by the geometric mean of growth and multiplication, and thus gives a doubling time of $\ln(2)/\kappa$.

### No self-replication

In the absence of self-replication, the two remaining processes are the spontaneous appearance and the removal of aggregates. The resulting system predicts a random independent number of aggregates in each synaptosome, i.e. a Poisson distribution. This can also be verified by following the above derivation with $\kappa = 0$.
